# Separation and detection of minimal length isomeric glycopeptide neoantigen epitopes centering GSTA region of MUC1 by LC-MS

**DOI:** 10.1101/617308

**Authors:** Dapeng Zhou, Kaijie Xiao, Zhixin Tian

**Affiliations:** School of Medicine, Tongji University, Shanghai 200092, China; School of Chemical Science and Engineering & Shanghai Key Laboratory of Chemical Assessment and Sustainability, Tongji University, Shanghai 200092, China

**Keywords:** MUC1, glycopeptide, mass spectrometry, electron transfer dissociation, neoantigen, epitope

## Abstract

MUC1 ranks No.2 on the list of targets for cancer immunotherapy. We previously reported monoclonal antibodies binding to glycopeptide neoantigen epitopes centering GSTA sequence of the highly glycosylated tandem repeat region of MUC1. Epitopes centering GSTA sequence are also predicted by NetMHC programs to bind to MHC molecules, although empirical data are lacking. Detecting isomeric MUC1 glycopeptide epitopes by mass spectrometry (MS) remains a technical challenge since antigenic epitopes are often shorter than 10 amino acids. MUC1 digests by Arg-C-specific endopeptidase clostripain could generate heterogenous icosapeptides, but isomeric 20-residue glycopetides could not be separated by liquid chromatography. In this study, we used pronase from *Streptomyces griseus*, which has no amino acid sequence preference for enzymatic cleavage sites, to digest a pair of synthetic glycopeptide isomers RPAPGST(Tn)APPAHG and RPAPGS(Tn)TAPPAHG, and analyzed the digests by LC-MS using electron transfer dissociation (ETD) and higher-energy collisional dissociation (HCD) methods. The results showed that glycopeptide isomers containing 8 to 11 amino acids could be efficiently generated by pronase digestion. Such glycopeptide isomers of minimal epitope lengths were clearly distinguished by characteristic MS/MS ion patterns and elution profiles of liquid chromatography. A glycopeptide library was generated which may serve as standards for measuring neoantigen epitopes centering GSTA sequence.

## Introduction

MUC1 is a type I membrane glycoprotein currently ranked as number 2 on the list of targets for cancer immunotherapy (1-2). Its extracellular portion contains 20 to 120 or more tandem repeats (TR) of 20-residue sequence (HGVTSAPDTRPAPGSTAPPA) with five potential O-linked glycosylation sites through N-acetylgalactosamine to serine and/or threonine residues. It is expressed at low levels at the apical surface of most healthy glandular epithelial cells, with normal pattern of O-glycosylation. In tumor settings MUC1 loses its polarity and normal pattern of glycosylation to expose GalNAc (Tn) residue.

We and others have dissected the abnormally glycosylated TR region as three antigenic motifs: PDTR, GSTA, and GVTS (2). The glycosylation of the three aforementioned peptide motifs may influence their binding to mAbs by several mechanisms. Identification of glycosylation site on above motifs is critical for dissecting the exact MUC1 antigenic epitopes for cancer diagnosis and therapy. For example, we have reported the expression of abnormally glycosylated GSTA neoantigen motif in non-small cell carcinoma cells (3-4).

MUC1 glycopeptides may also be processed by MHC Class I and/or MHC class II pathway, and serve as neoantigen epitopes for T cells. Barnea et al. reported the elution of MUC1-derived NLTISDVSV sequence from HLA-A2 molecules in breast cancer cell line MCF-7, although glycosylated NLTISDVSV has not been reported (5-6). Hanisch et al. reported the processing of a recombinant glycoprotein containing six MUC1 tandem repeats by mouse dendritic cells, that glycopeptides centering all PDTR, GSTA, and GVTS motifs could be generated by lysosomal protease digestion (7). Ninkovic et al. reported the loading of SAPDT(GalNAc)RPAPG by HLA-A2 molecule, that stimulated CTL cells (8). However, the direct detection of endogenous glycopeptides eluted from MHC molecules have not been reported.

Significant progresses have been made for glycopeptide analysis in past decade. Databases (9-10) and methods (11-31) for analyzing N-glycopeptides and O-glycopepitdes are accumulating, especially for site-specific glycosylation. Obviously, data containing both identities of glycan structures and glycosylation sites are most valuable for their functional analysis, compared to data containing glycan structures or glycosylation sites alone.

The accumulation of glycopeptide data benefit from most widely used MS ionization and ion activation techniques. Especially, electro-spray ionization (ESI) and electron transfer dissociation (ETD) greatly paved studies on O-linked glycosylation, allowing in-depth analysis of both O-glycan structures and the occupancy of O-glycosylation sites. Levery and Clausen group published systemic analysis of O-glycopeptides in trypsin-digested proteins from CHO cells (21). Zhang group recently published site-specific extraction of O-linked glycopeptides in trypsin-digested tissues (31). However, no data has been available for the TR region of mucin-1 protein yet due to the resistance of TR region to trypsin digestion. Muller et al. used Arg-C-specific endopeptidase clostripain to digest human milk MUC1, which generated heterogenous icosapeptides starting with the PAP sequence. O-glycosylation sites were localized by post-source decay matrix-assisted laser desorption ionization mass spectrometry or by solid phase Edman degradation (32). Sihlbom et al. used LC-MS with electron-capture dissociation (ECD) fragmentation method to analyze clostripain-digested icosapeptides, and could localize the O-glycoylation sites in heterogenous icosapeptides (33). However, the separation of isomeric icosapeptides could not be achieved by liquid chromatography.

In this study, we tested the efficacy of pronase from *Streptomyces griseus* in digesting synthetic MUC1 glycopeptides, and generated short glycopeptides library containing 8 to 11 amino acids.

## Methods

### Synthesis of RPAPGST(Tn)APPAHG and RPAPGS(Tn)TAPPAHG

Antigenic epitopes for antibody binding are mostly shorter than 10 amino acids. We chose to use two synthetic 13-residue glycopeptides, RPAPGST(Tn)APPAHG and RPAPGS(Tn)TAPPAHG, as the starting material for generating glycopeptides shorter than 10 amino acids. The chemical synthesis of glycopeptides was as described (3) using fluorenylmethyloxycarbonyl (Fmoc)-protected amino acids as the building blocks. The purity of synthetic glycopeptides were examined by reversed-phase HPLC and MS determination of molecular masses.

### Pronase digestion of MUC1 glycopeptides

Glycopeptides were digested by pronase from *Streptomyces griseus* (Roche Diagnostics, Germany) according to the manufacturer’s protocol. In brief, 5 µg glycopeptides were digested in 100 µL Tris buffer (50 mM, pH 7.6) with 5 mM CaCl_2_ and 10 mg/mL pronase at 50 °C for 12 hours. The pronase was inactivated by heating to 100 °C for 5 minutes. The digests were desalted by a home-made C18 reverse chromatography column, before analyzed by mass spectrometry.

### LC-MS/MS analysis of pronase digests of RPAPGST(Tn)APPAHG and RPAPGS(Tn)TAPPAHG

The pronase digests of both RPAPGST(Tn)APPAHG and RPAPGS(Tn)TAPPAHG were analyzed on an Orbitrap Fusion Lumos MS (Thermo Scientific, San Jose, CA, USA) coupled with a nano-ESI source and a Dionex Ultimate 3000 RSLCnano HPLC system.

A C18 (Agilent ZORBAX 300SB, 5 μm, 300 Å) pre-column (360 μm o.d. × 200 μm i.d., 7 cm long) was used for sample loading. Chromatographic separation was performed on a 75-cm-long analytical column (360 μm o.d. × 75 μm i.d.) packed with the same C18 particles with the pre-column; buffer A is composed of 99.9% H_2_O and 0.1% FA, and buffer B is composed of 99.9% ACN and 0.1% FA. The flow rate of the mobile phase was 300 nL/min with a multi-step gradient starting with 4% B: 8% B, 4 min; 30% B, 49 min; 100% B, 4 min; 100% B, 3 min.

MS spectra were acquired as follows: mass resolution 60 k; m/z range 375-1800, max ion injection time 50 ms, automatic gain control (AGC) target 4e5, microscans 1, RF lens 40%. MS/MS spectra were acquired with the following settings: data-dependent mode, cycle time 3 s, isolation width 1.2 Th, first mass 100, ETD activation time 200 ms, ETD reagent target 2e5, ETD reagent injection time 200 ms, supplemental activation 35%, mass resolution 30 k, max ion injection time 200 ms, AGC target 5e4, microscans 1, dynamic exclusion 30 s, included charge states 2-7. The ESI conditions were as follows: spray voltage 2.2 kV, capillary temperature 320 °C.

### Database search and peptide identification

Database search and peptide identification from the LC-MS/MS datasets of the full-length synthetic peptides and their pronase digests were carried out using ProteinGoggle; the detailed interpretation steps have been published elsewhere (34-39). Briefly, a customized peptide database was created for the full-length RPAPGS(Tn)TAPPAHG and RPAPGST(Tn)APPAHG, as well as their pronase digests with GalNAc as a dynamic modification on either S or T, containing theoretical isotopic envelopes information of both the precursor and fragment ions (a, b, c, x, y, and z). Matched precursor and fragment ions were searched with the following IPACO/IPMD/IPAD parameters: 40%/15ppm/100%, 20%/15 ppm/50%.

## Results

### Synthesis and detection of GalNAc-modified MUC1 antigen epitopes centering GSTA motif

The shortest glycopeptides of MUC1 tandem repeat region that have been detected in the past were icosapeptides (20-residue glycopeptides) prepared from MUC1 protein digests by Arg-C-specific endopeptidase clostripain (32-33). However, antibody-binding epitopes are often shorter than 10 amino acids. We previously tested a 13-residue glycopeptide, RPAPGS(Tn)TAPPAHG, which contains antibody binding sites for 16A (3), and synthesized its isomer RPAPGST(Tn)APPAHG.

The glycopeptide RPAPGS(Tn)TAPPAHG was confidently detected as shown in the MS spectrum (Figure 1A) with IPAD value within 5% and IPMD value within 5 ppm. 18 matched fragment ions were observed in the MS/MS spectrum (Figure 1B, *=GalNAc), among which 8 contain GalNAc (as shown in the red box). We detected c6* (*m/z* 786.40936) and z7 (*m/z* 634.30573), the two theoretical glycosite-determining fragment ions.

**Figure 1.**
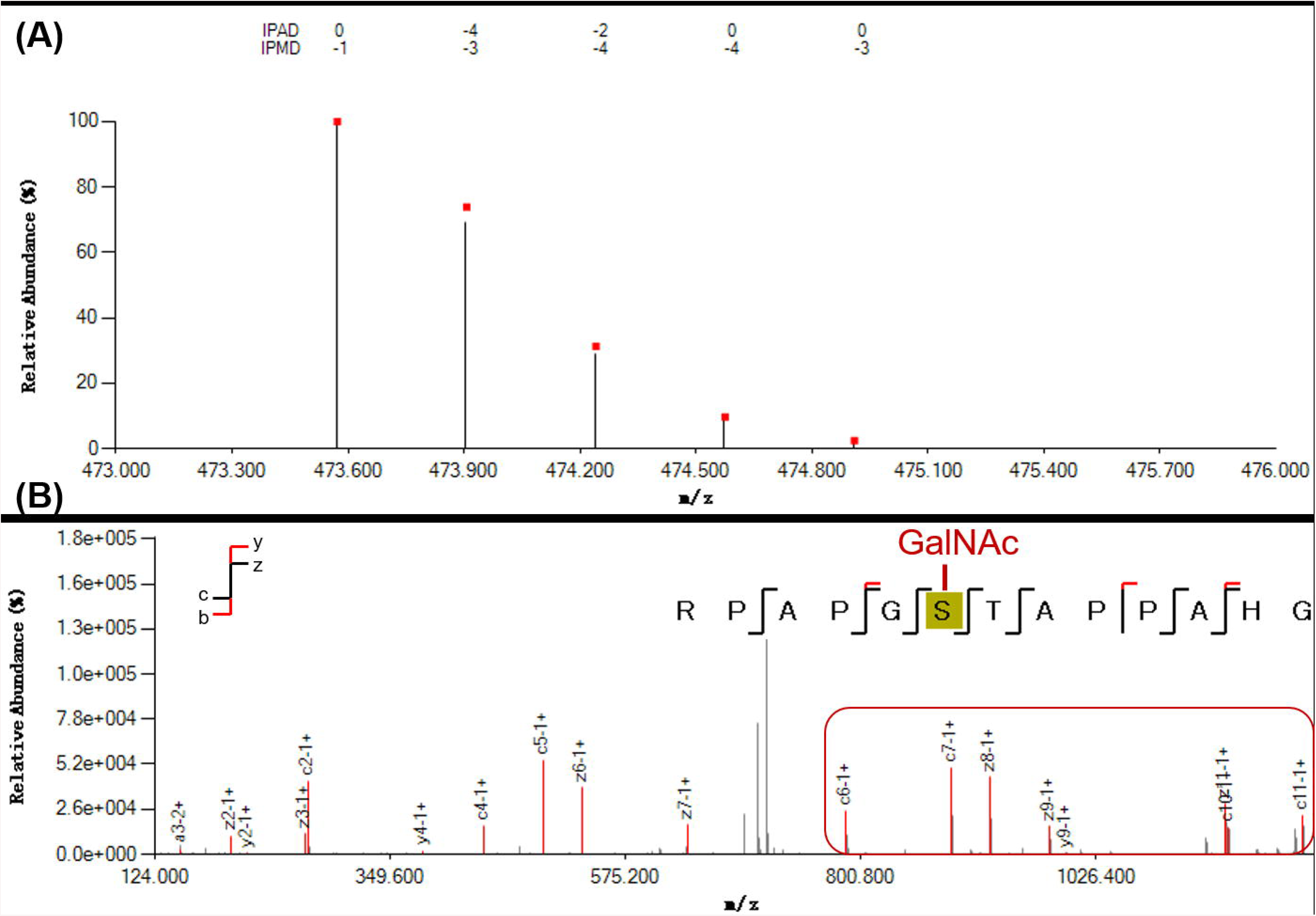
MS and MS/MS spectra for RPAPGS(Tn)TAPPAHG. (A) The isotopic envelope fingerprinting map of the precursor ion. Red squares indicate theoretical mass spectra, and black bars indicate experimental mass spectra. (B) The annotated MS/MS spectrum with the graphical fragmentation map; the fragment ions containing GalNAc are marked in red box. The glyco-site determining fragment ions are c6-1^+^* (*=GalNAc, *m/z* 786.40936) and z7-1+ (*m/z* 634.30573). Red bars indicate that the experimental mass spectra match with the theoretical mass spectra.

We also detected the glycopeptide RPAPGST(Tn)APPAHG (Figure 2A) with IPAD and IPMD values lower than 4% and 4 ppm, respectively. Also observed in this peptide included 18 matched fragment ions in the MS/MS spectrum (Figure 2B, *=GalNAc), among which 9 contain GalNAc (as shown in the read box) including c6 (*m/z* 583.33026) and z7* (*m/z* 837.38580).

**Figure 2.**
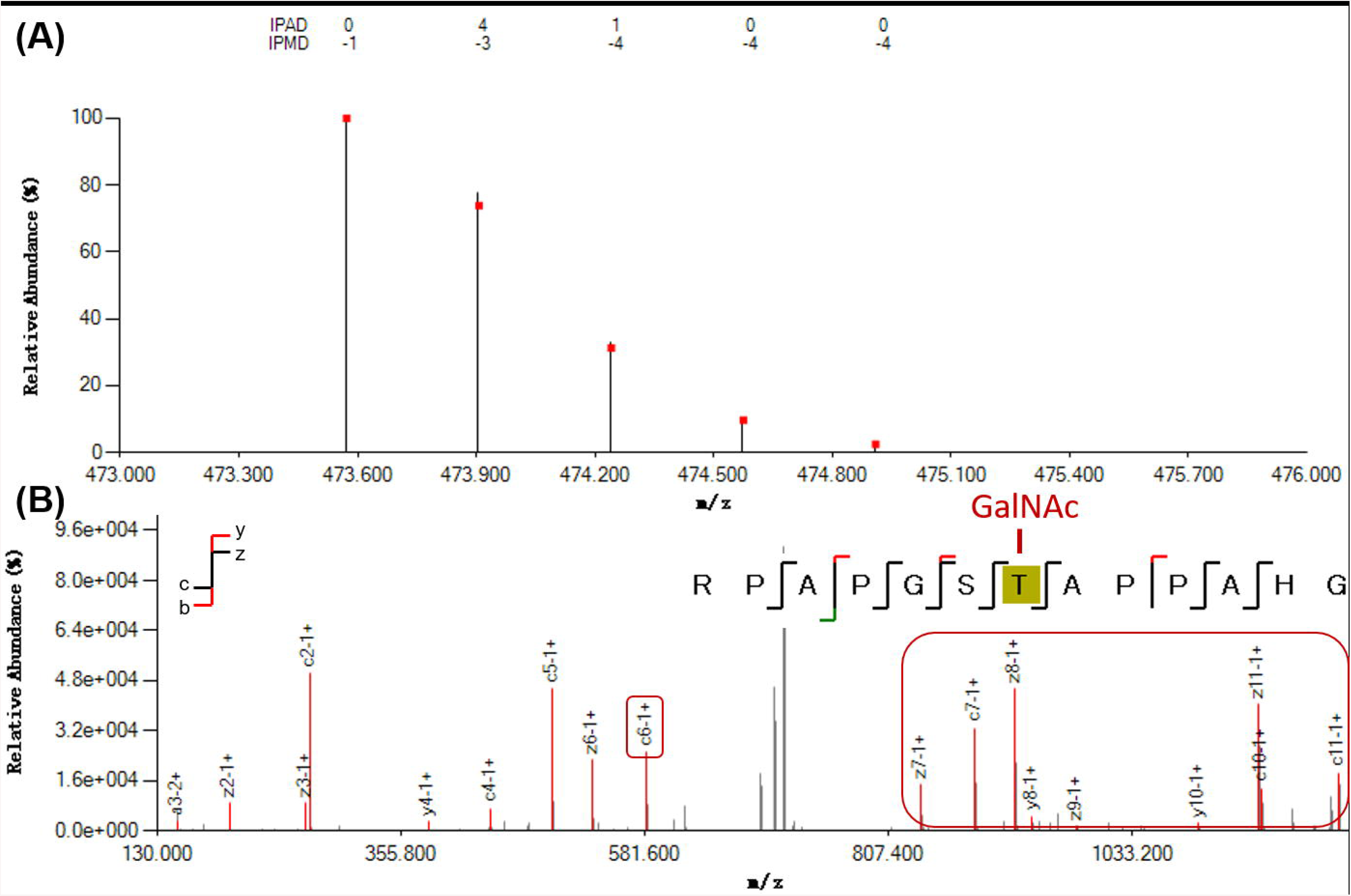
MS and MS/MS profiles for RPAPGST(Tn)APPAHG. (A) The isotopic envelope fingerprinting map of the precursor ion. Red squares indicate theoretical mass spectra, and black bars indicate experimental mass spectra. (B) The annotated MS/MS spectrum with the graphical fragmentation map; the fragment ions containing GalNAc are marked in red box. The glyco-site determining ions are c6-1+ (*m/z* 583.33026) and z7-1+* (*=GalNAc, *m/z* 837.38580). Red bars indicate that the experimental mass spectra match with the theoretical mass spectra.

### A MS/MS library of MUC1 antigen epitope glycopeptides with 8 to 11 amino acids

We further analyzed the pronase digests of chemically synthesized 13-residue glycopeptides. In contrast to trypsin, or Arg-C-specific endopeptidase clostripain, which are site-specific, the pronase cleaves peptide-bonds without sequence-specificity. The cleaved glycopeptides could be detected as listed in Table 1.

**Table 1.** Short glycopeptides from pronase digests of RPAPGS(Tn)TAPPAHG and RPAPGST(Tn)APPAHG

|  | Amino acid<br>sequence | Sequence length | Site-determining<br>product ions | MS-MS<br>profile | Retention time<br>(min) |
| --- | --- | --- | --- | --- | --- |
| -GS(Tn)TA- | RPAPGS(Tn)TAPPA | 11 | c6-1<sup>+</sup>* (*m/z* 786.41089) | Figure S1 | 15.24 |
|  | RPAPGS(Tn)TAPP | 10 | c6-1<sup>+</sup>* (*m/z* 786.40985) | Figure S2 | 11.11 |
|  | APGS(Tn)TAPPA | 9 | c4-1<sup>+</sup>* (*m/z* 533.25671) | Figure S3 | 12.17 |
| -GST(Tn)A- | RPAPGST(Tn)APPA | 11 | c6-1<sup>+</sup> (*m/z* 583.33124) | Figure S4 | 11.15 |
|  | RPAPGST(Tn)APP | 10 | c6-1<sup>+</sup> (*m/z* 583.33008) | Figure S5 | 11.10 |
|  | APGST(Tn)APPA | 9 | b4-1<sup>+</sup> (*m/z* 313.15085) | Figure S6 | 12.00 |
|  |  |  | c4-1<sup>+</sup> (*m/z* 330.17761) |  |  |
|  |  |  | y5-1<sup>+</sup>* (*m/z* 659.32452) |  |  |
|  | APGST(Tn)APP | 8 | b4-1<sup>+</sup> (*m/z* 313.15097) | Figure S7 | 12.86 |
|  |  |  | c4-1<sup>+</sup> (*m/z* 330.17856) |  |  |

For the pronase digests of RPAPGS(Tn)TAPPAHG, the 11-, 10- and 9-residue glycopeptides were successfully identified with unambiguous glycosite localization (Table 1). The 11-residue was observed with 9 matched fragment ions (Figure S1); b6*/c6* and y5/z5 are the four theoretical glycosite-determining fragment ions, and c6-1+* was actually observed. The 10-residue was observed with 11 matched fragment ions (Figure S2); b6*/c6* and y4/z4 are the four theoretical glycosite-determining fragment ions, and c6-1+* was actually observed. The 9-residue was observed with 10 matched fragment ions (Figure S3); b4*/c4* and y5/z5 are the four theoretical glycosite-determining fragment ions, and c4-1+* was actually observed.

For the pronase digests of RPAPGST(Tn)APPAHG, the 11-, 10-, 9- and 8-residue glycopeptides were successfully identified with unambiguous glycosite localization (Table 1) as well. The 11-residue was observed with 6 matched fragment ions (Figure S4); b6/c6 and y5*/z5* are the four theoretical glycosite-determining fragment ions, and c6-1+ was actually observed. The 10-residue was observed with 5 matched fragment ions (Figure S5); b6/c6 and y4*/z4* are the four theoretical glycosite-determining fragment ions, and c6-1+ was actually observed. The 9-residue was observed with 14 matched fragment ions (Figure S6); b4/c4 and y5*/z5* are the four theoretical glycosite-determining fragment ions, and b4-1+, c4-1+ and y5-1+* were actually observed. The 8-residue was observed with 8 matched fragment ions (Figure S7); b4/c4 and y4*/z4* are the four theoretical glycosite-determining fragment ions, and b4-1+ and c4-1+ were actually observed.

## Discussion

### Feasibility to distinguish MUC1 glycopeptide isomers shorter than 10 amino acids by LC-MS methods

Our data showed that it is feasible to distinguish glycosite isomers of glycopeptides by LC-MS/MS method. The signature glycosite-determining fragment ions together with differential retention time can clearly separate the paired glycosite isomers of RPAPGS(Tn)TAPPA vs. RPAPGST(Tn)APPA, RPAPGS(Tn)TAPP vs. RPAPGST(Tn)APP, and APGS(Tn)TAPPA vs. APGST(Tn)APPA (Table 1).

### Feasibility to generate short MUC1 glycopeptide libraries by pronase

The TR region of MUC1 is resistant to trypsin digestion. Previous attempts to digest MUC1 by Arg-C-specific endopeptidase clostripain could generate heterogenous icosapeptides, but isomeric 20-residue glycopetides could not be separated by liquid chromatography (32-33). To generate minimal epitope length glycopeptides for LC-MS identification, we digested the 13-residue glycopeptides by pronase from *Streptomyces griseus*. We could generate 9-, 10- and 11-residue short glycopeptide isomers which could be clearly identified by LC-MS/MS.

Among the pronase digests of RPAPGST(Tn)APPAHG, the 7-residue APGS(Tn)TAP was well observed in the MS spectrum (Figure S8A), and 8 matched fragment ions were observed in its EThcD MS/MS spectrum (Figure S8B); b4*/c4* and y3/z3 are the four theoretical glycosite-determining fragment ions, but none of them was observed. Among the pronase digests of RPAPGST(Tn)APPAHG, the 7-residue APGST(Tn)AP was well observed in the MS spectrum (Figure S9A), and 7 matched fragment ions were observed in its EThcD MS/MS spectrum (Figure S9B);, b4/c4 and y3*/z3* are the four theoretical glycosite-determining fragment ions, but none of them was observed.

The 12-residue digest of both RPAPGS(Tn)TAPPAHG and RPAPGST(Tn)APPAHG, and the 8-residue digest of the former were not identified in the current study; most likely, they are not abundantly produced during pronase digestion. Alternatively, it might be due to their inefficient ionization.

In summary, our data established the LC-MS/MS identities of a clinically-relevant MUC1 glycopeptide neoantigen epitope centering GSTA motif. A library of short MUC1 glycopeptides centered on GSTA motif was created, which is a critical step for analysis of such antigen epitopes in real biological samples.

## Abbreviations list

TR: tandem repeat;
Tn: GalNAc;
mAb: monoclonal antibody;
MS: mass spectrometry;
ESI: electro-spray ionization;
ETD: electron transfer dissociation;
HCD: higher-energy collisional dissociation;
IPACO: isotopic peak abundance cutoff;
IPMD: isotopic peak *m/z* deviation;
IPAD: isotopic peak abundance deviation.

## Acknowledgements

We thank Drs. Wei Zhu and Wenzhang Chen at Proteomics Core Facility of ShanghaiTech University, Dr. Hangxin Cheng at Thermo Shanghai Service Center for technical assistance.

## Author contribution statement

DZ designed this study. DZ, KX, and ZT contributed to the collection, analysis and interpretation of data. DZ and ZT wrote the manuscript. All authors read and approved the final manuscript.

