## Supplementary information for "Separation and detection of minimal length isomeric glycopeptide neoantigen epitopes centering GSTA region of MUC1 by LC-MS"

**Figure S1.** MS and MS/MS profiles for RPAPGS(Tn)TAPPA (11 amino acids). (A) The isotopic envelope fingerprinting map of the precursor ion; (B) The annotated MS/MS spectrum with the graphical fragmentation map; the fragment ions containing GalNAc are marked in red boxes. The glycosite determining fragment ion is c6-1^+^*( *=GalNAc).

**Figure S2.** MS and MS/MS profiles for RPAPGS(Tn)TAPP (10 amino acids). (A) The isotopic envelope fingerprinting map of the precursor ion; (B) The annotated MS/MS spectrum with the graphical fragmentation map; the fragment ions containing GalNAc are marked in red boxes. The glycosite determining fragment ion is c6-1^+^*(*=GalNAc).

**Figure S3.** MS and MS/MS profiles for APGS(Tn)TAPPA (9 amino acids). (A) The isotopic envelope fingerprinting map of the precursor ion; (B) The annotated MS/MS spectrum with the graphical fragmentation map; the fragment ions containing GalNAc are marked in red boxes. The glycosite determining fragment ion is c4-1^+^*(*=GalNAc).

**Figure S4.** MS and MS/MS profiles for RPAPGST(Tn)APPA (11 amino acids). (A) The isotopic envelope fingerprinting map of the precursor ion; (B) The annotated MS/MS spectrum with the graphical fragmentation map; the fragment ions containing GalNAc are marked in red boxes. The glycosite determining fragment ion is c6-1^+^.

**Figure S5.** MS and MS/MS profiles for RPAPGST(Tn)APP (10 amino acids). (A) The isotopic envelope fingerprinting map of the precursor ion; (B) The annotated MS/MS spectrum with the graphical fragmentation map; the fragment ions containing GalNAc are marked in red boxes. The glycosite determining fragment ion is c6-1^+^.

**Figure S6.** MS and MS/MS profiles for APGST(Tn)APPA (9 amino acids). (A) The isotopic envelope fingerprinting map of the precursor ion; (B) The annotated MS/MS spectrum with the graphical fragmentation map; the fragment ions containing GalNAc are marked in red boxes. The glycosite determining fragment ions are b4-1+, c4-1+ and y5-1^+^*(*=GalNAc).

**Figure S7.** MS and MS/MS profiles for APGST(Tn)APP (8 amino acids). (A) The isotopic envelope fingerprinting map of the precursor ion; (B) The annotated MS/MS spectrum with the graphical fragmentation map. The glycosite determining fragment ions are b4-1+ and c4-1+.

**Figure S8.** MS and MS/MS profiles for APGS(Tn)TAP (7 amino acids). (A) The isotopic envelope fingerprinting map of the precursor ion; (B) The annotated MS/MS spectrum with the graphical fragmentation map. No glycosite determining fragment ion or ion pair is observed.

**Figure S9.** MS and MS/MS profiles for APGST(Tn)AP (7 amino acids). (A) The isotopic envelope fingerprinting map of the precursor ion; (B) The annotated MS/MS spectrum with the graphical fragmentation map. No glycosite determining fragment ion or ion pair is observed.
