## Supplementary figures and images for "Separation and detection of minimal length isomeric glycopeptide neoantigen epitopes centering GSTA region of MUC1 by LC-MS"

### Figure S1

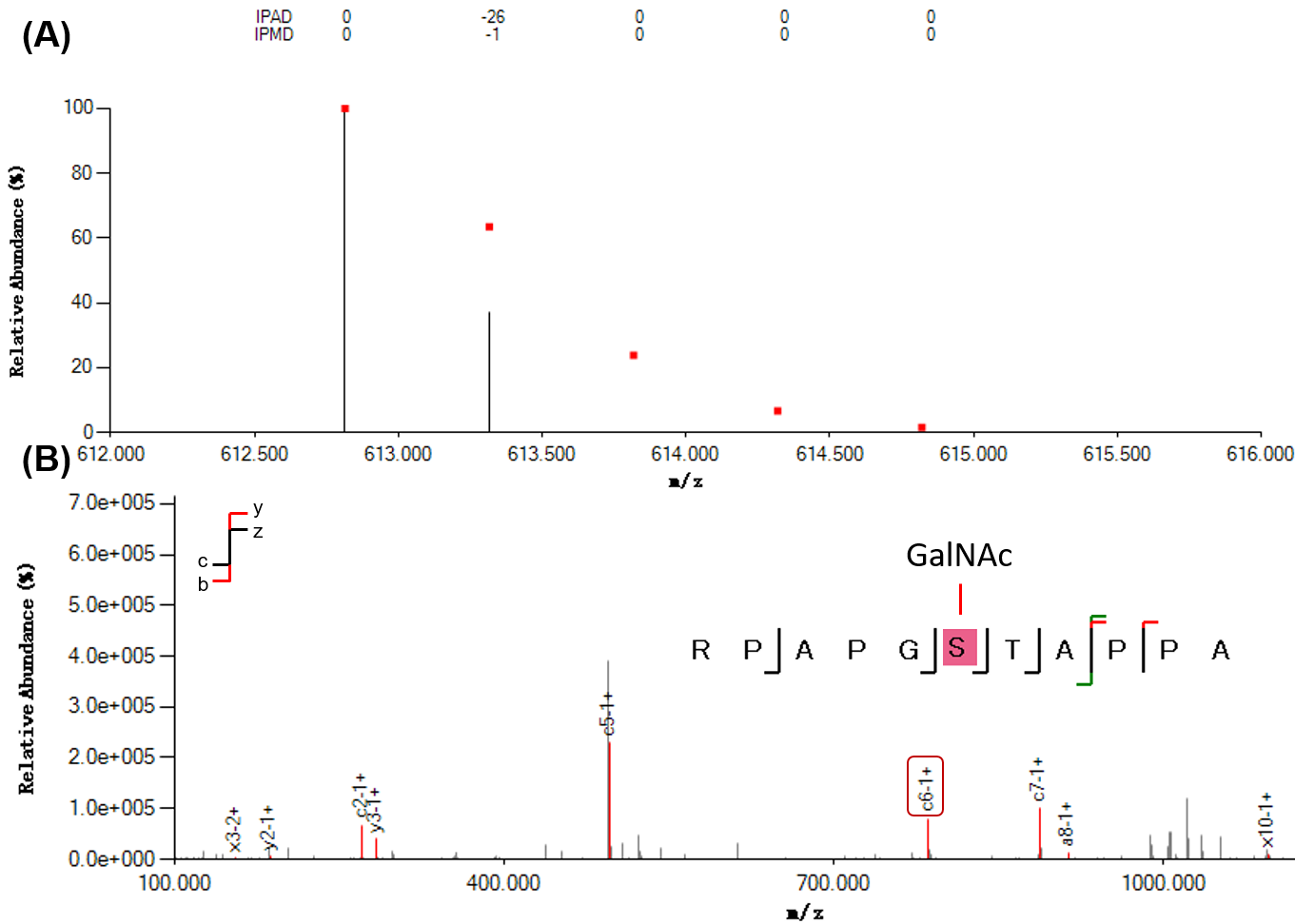

### Figure S2

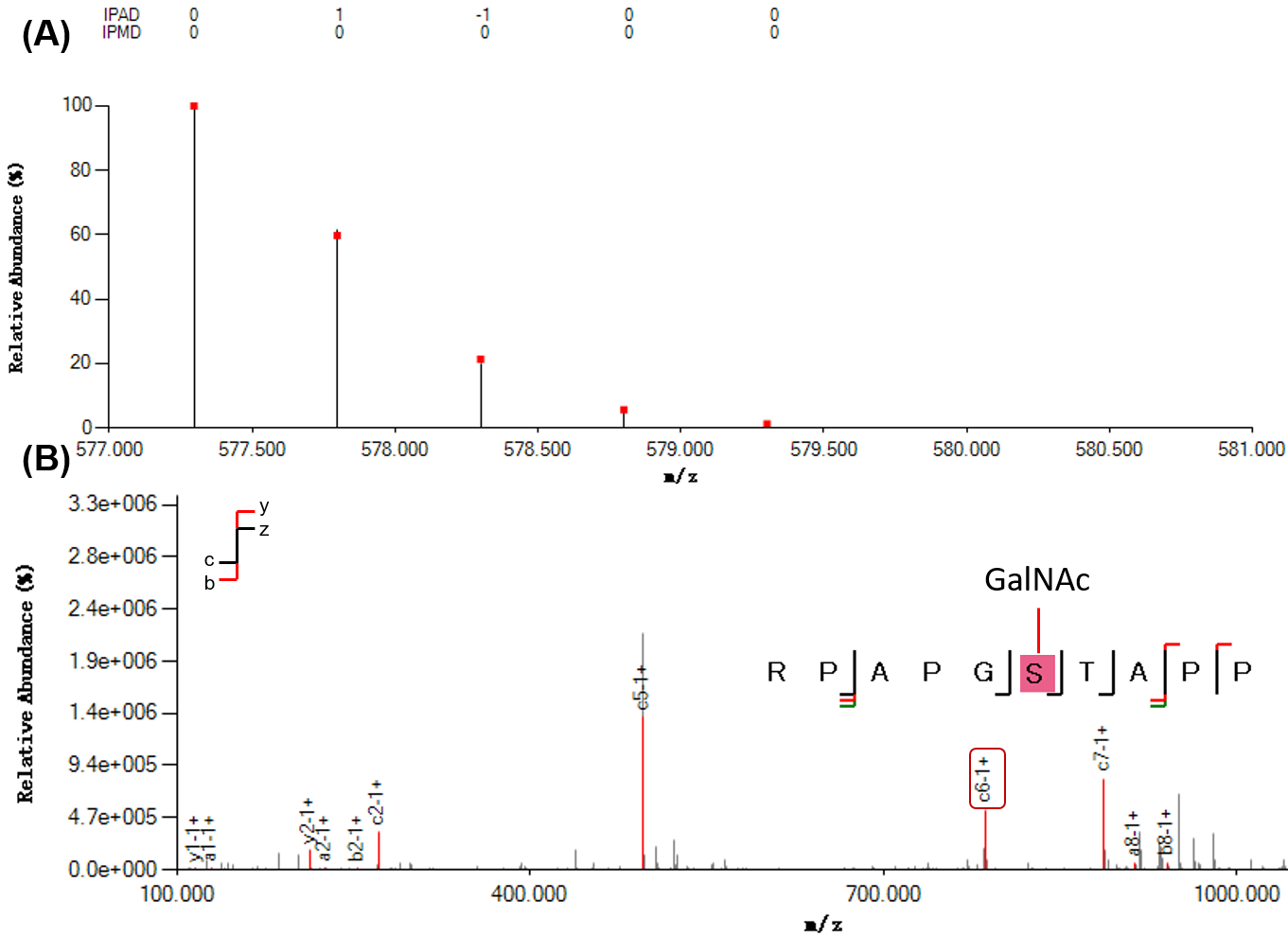

### Figure S3

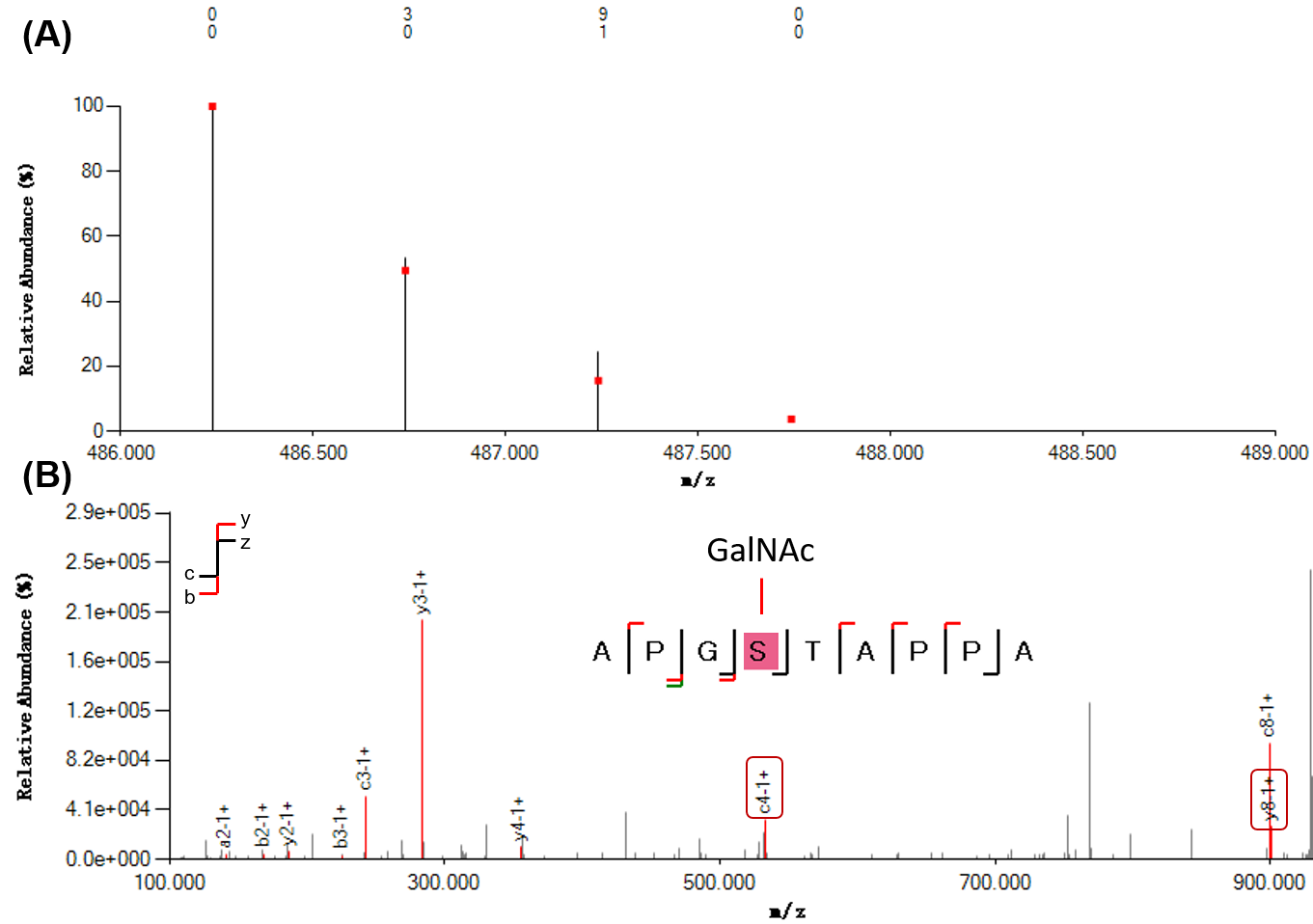

### Figure S4

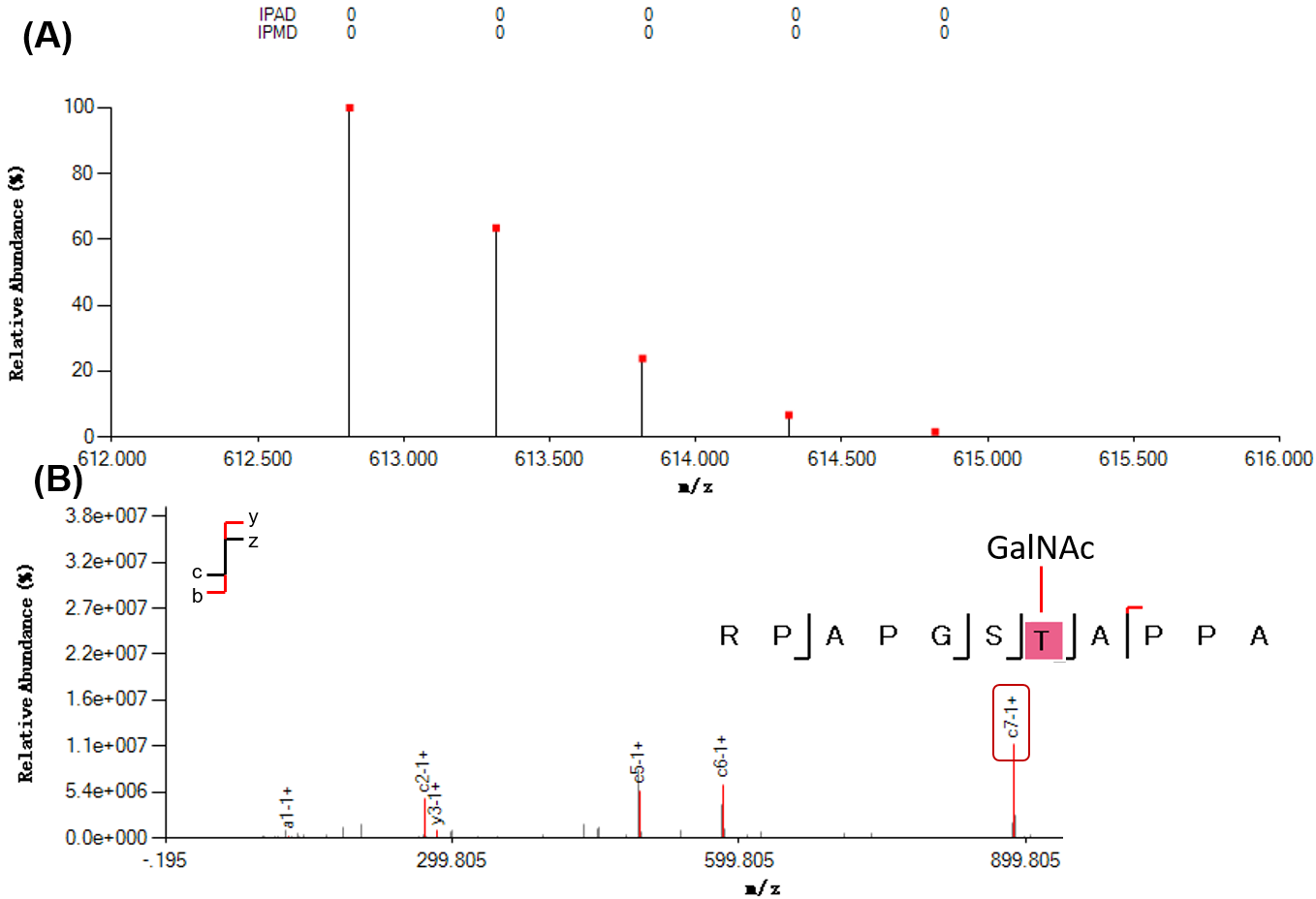

### Figure S5

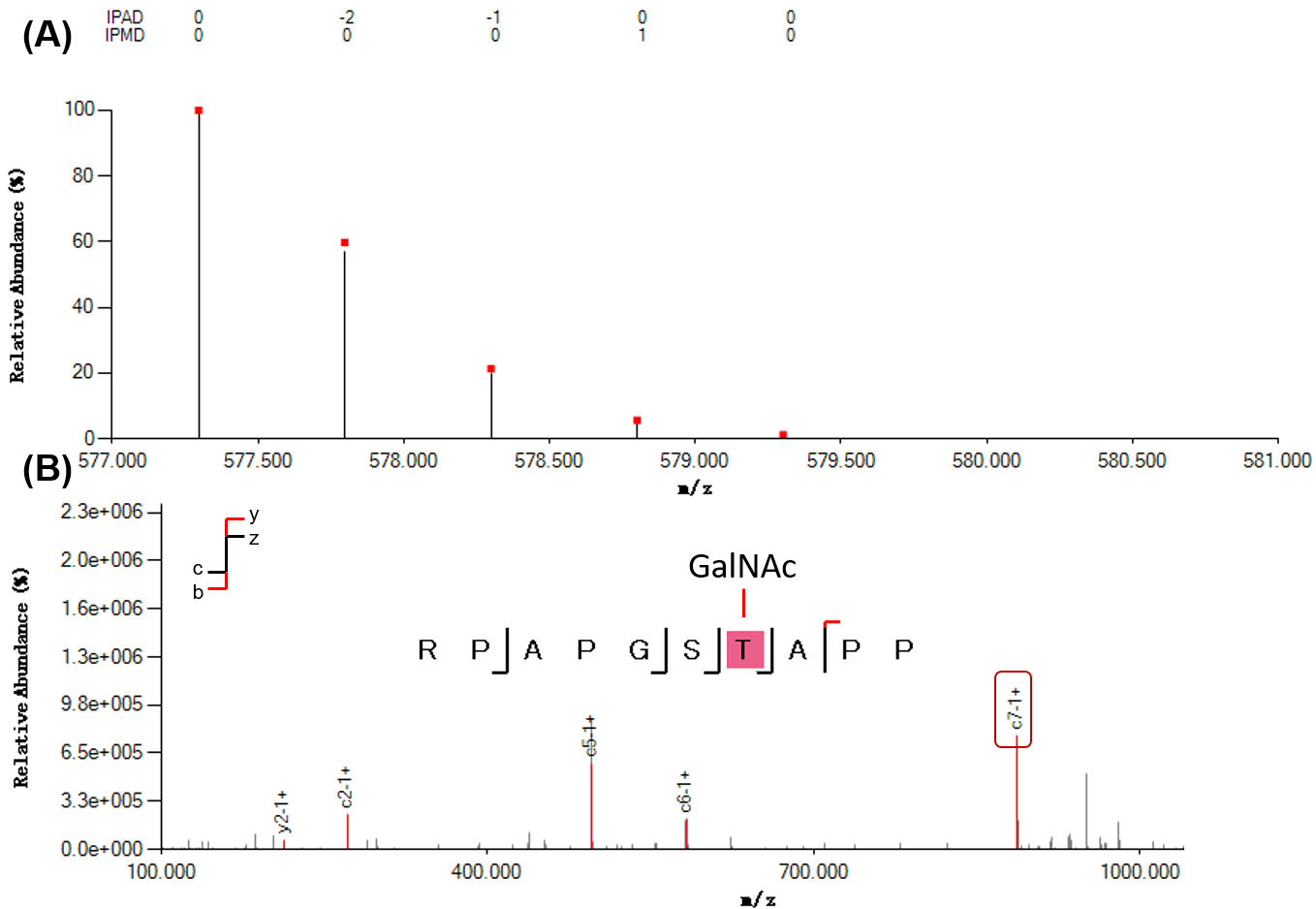

### Figure S6

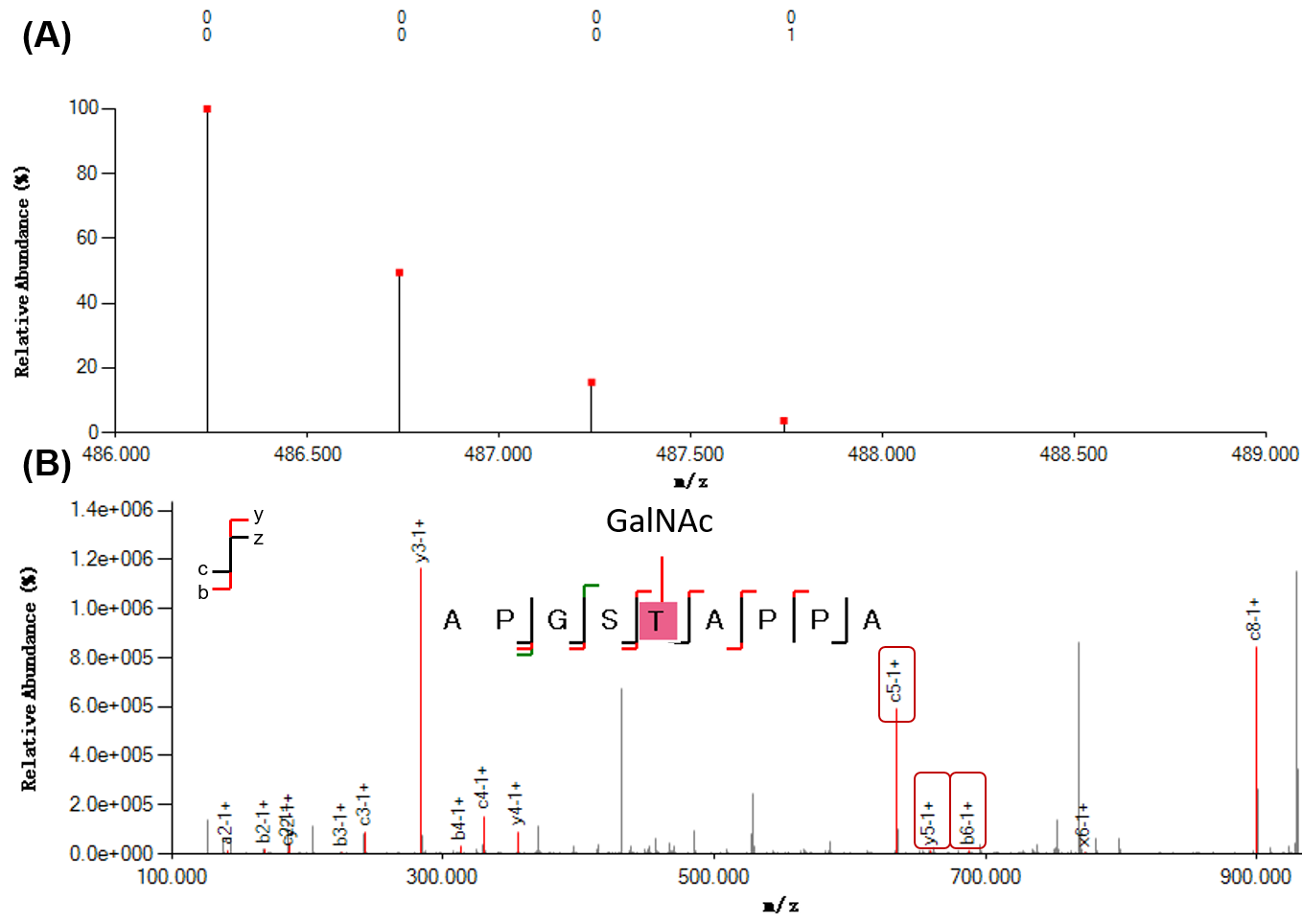

### Figure S7

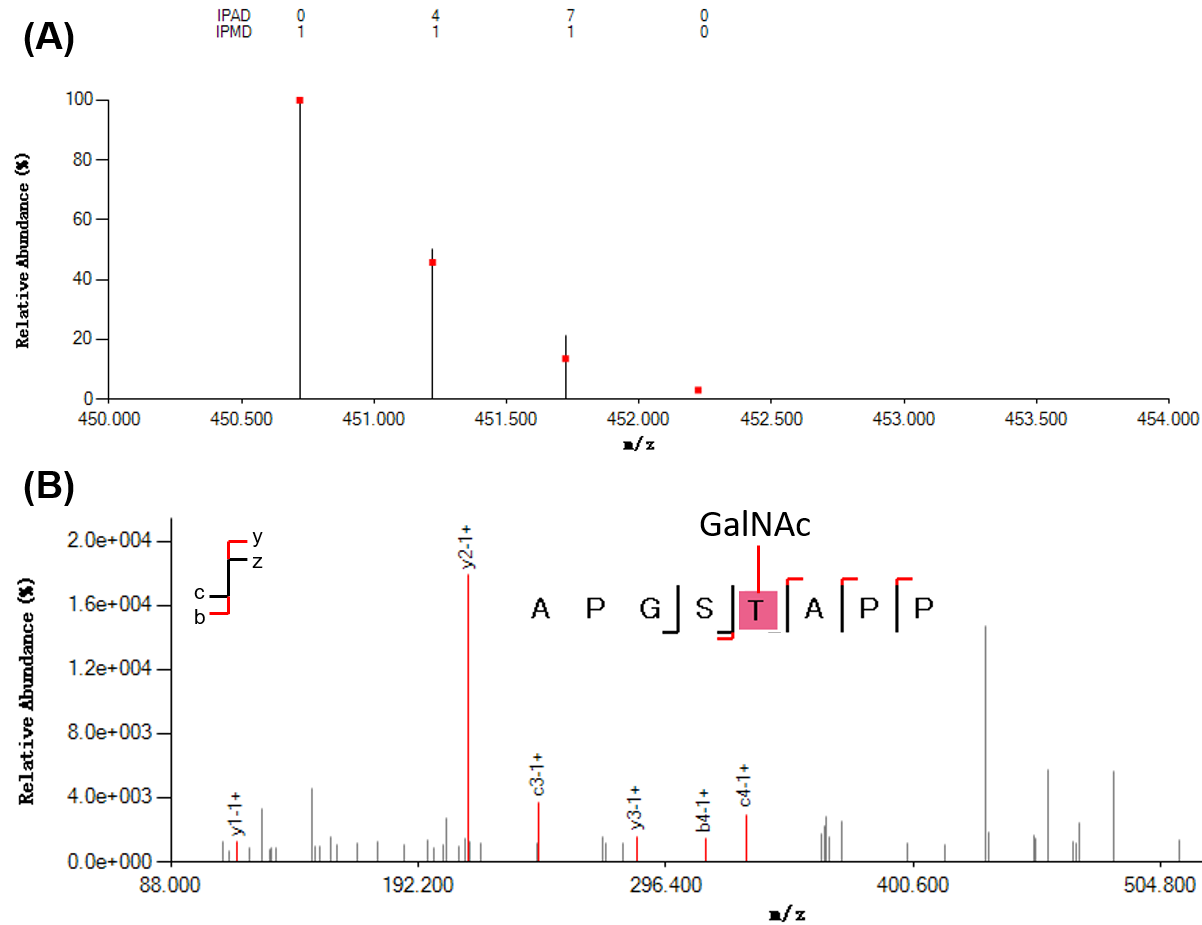

### Figure S8

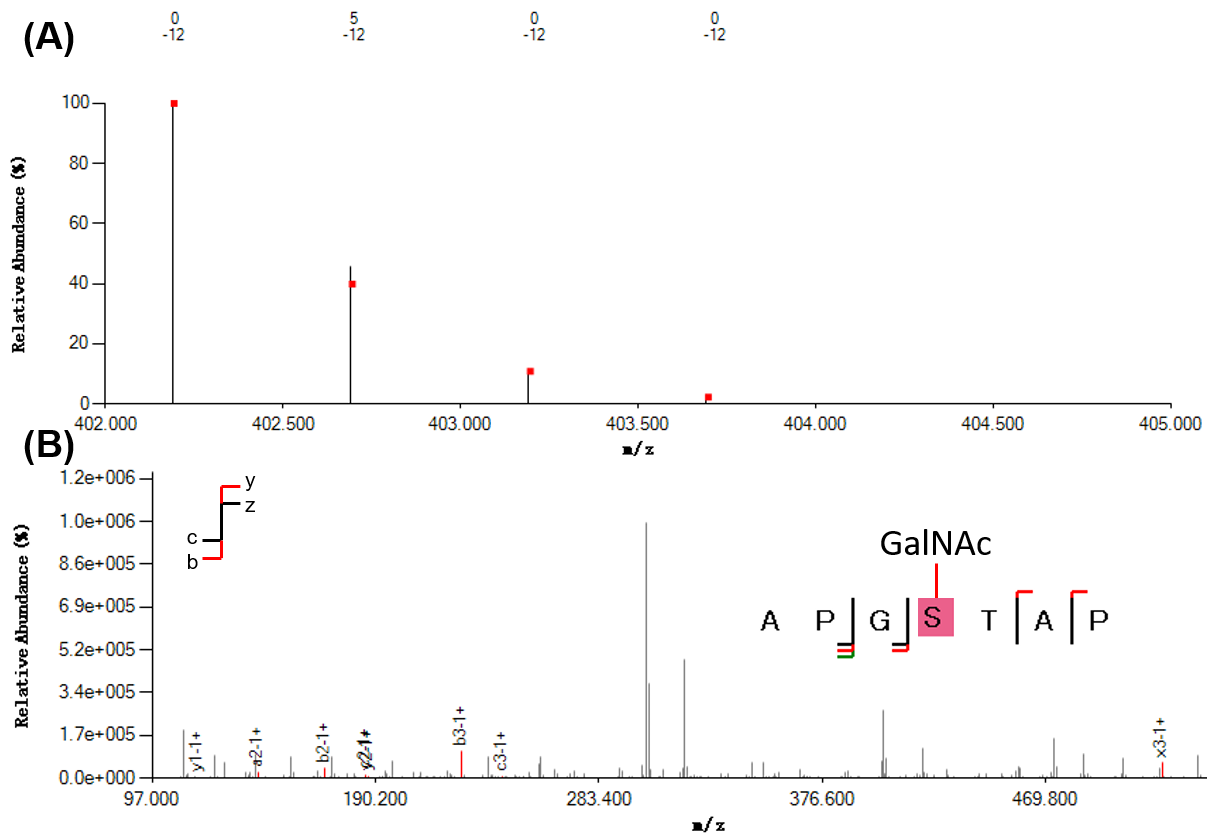

### Figure S9

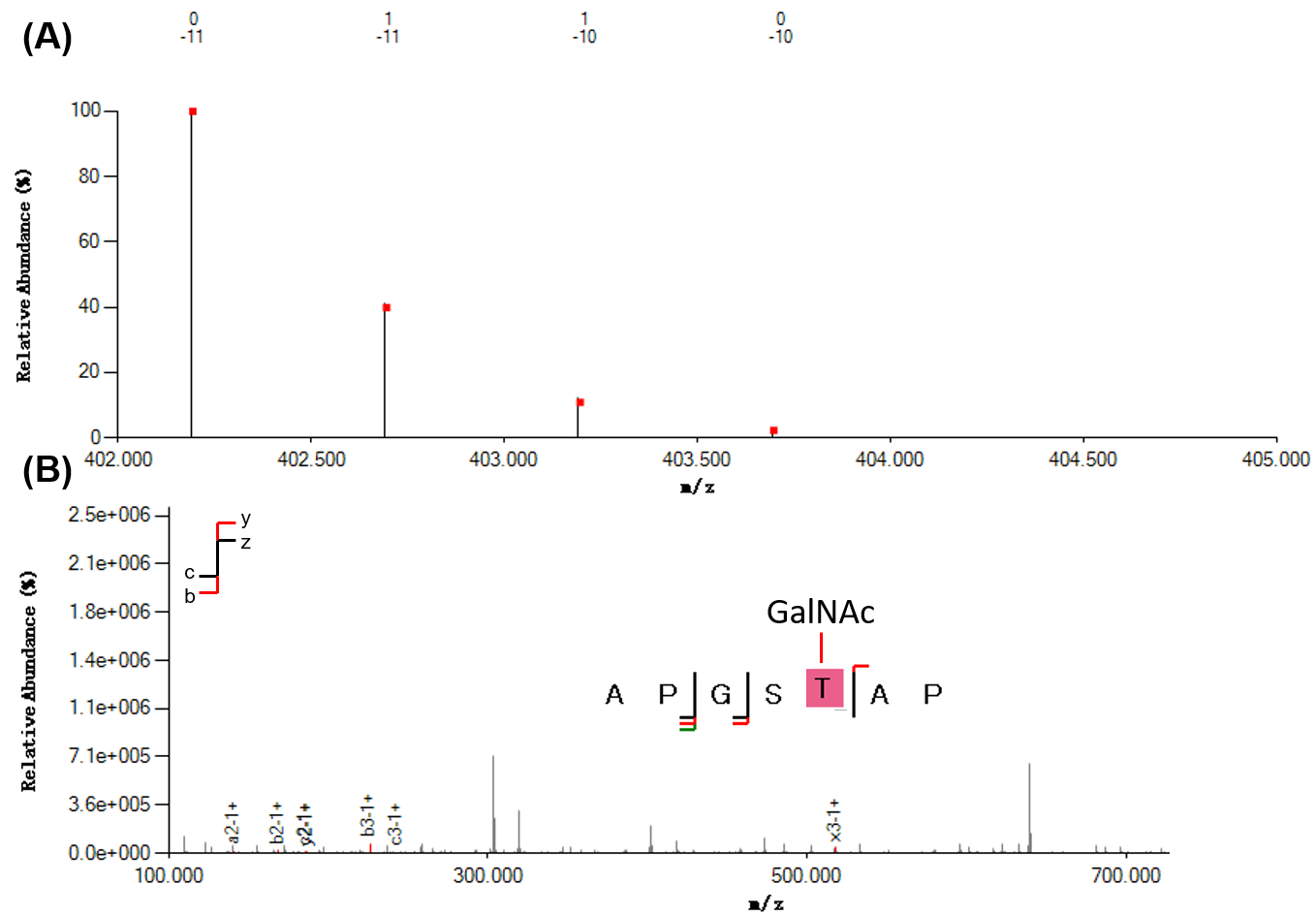
